# Spatiotemporal dynamics of alphaherpesviral latency and reactivation in the murine central nervous system

**DOI:** 10.1101/2025.09.11.675502

**Authors:** Viktoria Korff, Issam El-Debs, Barbara G. Klupp, Jens P. Teifke, Thomas C. Mettenleiter, Julia Sehl-Ewert

## Abstract

Alphaherpesviruses, including Herpes Simplex Virus 1 (HSV-1) and Pseudorabies Virus (PrV), establish lifelong latency in the nervous system and can cause recurrent disease. While latency has classically been attributed to peripheral sensory ganglia, accumulating evidence indicates that the central nervous system (CNS) also act as a relevant reservoir for viral latency and reactivation. Here, we investigated the CNS as a site of latency using the attenuated mutant PrV-ΔUL21/US3Δkin which reproduces key features of herpes simplex encephalitis (HSE) in female CD1 mice. We mapped brain regions permissive to alphaherpesviral latency and analyzed the temporal dynamics of viral transcription, histopathology, and host clinical and immune responses. Following intranasal inoculation, mice were analyzed at 11 to 14, 21, 28, 42, 105, and 190 days post infection (dpi). To assess the potential of reactivation, a subset received cyclophosphamide/dexamethasone at 170 dpi. Viral transcripts were detected by RNAscope™ in situ hybridization and RT-qPCR targeting the lytic gene UL19 and the latency-associated transcript (LAT). Histopathological analyses included hematoxylin and eosin (H&E) staining and immunohistochemistry for CD3, Iba1, GFAP, and cleaved caspase-3. Major capsid protein (UL19) expression displayed marked regional and temporal heterogeneity, with prominent signals in mesiotemporal structures (piriform cortex, hippocampus, entorhinal cortex), coinciding with pronounced T-cell infiltration. LAT expression remained overall low, with a transient peak during the acute infection (11–14 dpi). RT-qPCR confirmed high viral transcript levels for both UL19 and LAT in mesiotemporal regions during early infection, while LAT expression returned to baseline levels thereafter. Histopathology demonstrated a transition from acute necrotizing meningoencephalitis to prolonged or recurrent low-grade inflammation, accompanied by glial activation and localized apoptosis. Notably, UL19 expression strongly correlated with CD3^+^ T-cell infiltration, particularly at 42 dpi. These findings define the spatiotemporal interplay between viral transcriptional activity and neuroinflammation and identify selected CNS regions as reservoirs for latent or recurrent alphaherpesvirus infection.

**Author Summary:** Alphaherpesviruses are pathogens that not only cause acute disease but also establish lifelong latency in the nervous system. Under certain conditions, they can reactivate and trigger recurrent disease. Using a genetically modified pseudorabies virus in mice, we mapped how latent and lytic phases occur within the brain over several months. By combining molecular, histological, and clinical analyses, we show that specific brain regions act as long-term viral reservoirs, where signs of inflammation and viral activity can reappear long after the initial infection. These findings provide new insights into how alphaherpesviruses persist in the central nervous system and suggest that recurrent or subclinical reactivation may contribute to long-term neurological complications, including those resembling human herpes simplex encephalitis.

## Introduction

Alphaherpesviruses such as Herpes Simplex Virus Type 1 (HSV-1) and Pseudorabies Virus (PrV) are neurotropic DNA viruses capable of establishing lifelong latency within the nervous system (1). Following primary mucosal infection, virions enter peripheral sensory neurons and undergo retrograde axonal transport to peripheral and autonomic ganglia, particularly the trigeminal ganglion (TG) (2, 3). Here, the viral genome persists in a latent, transcriptionally restricted state (4, 1, 5–8).

Productive infection is characterized by a temporally regulated cascade of immediate-early, early, and late gene expression (9, 10). In contrast, latency is defined by episomal genome maintenance in the absence of infectious virus production, with transcription largely restricted to the latency-associated transcripts (LATs). In PrV, these include an unstable 8.4 kb transcript, the large-latency transcript (LLT), a stable intron, eleven micro-RNAs, two small noncoding RNAs, and three transcripts antisense to the major LAT (10–12). LAT and their encoded micro RNAs synergistically inhibit apoptosis (13, 14), and suppress viral replication (15, 16). Establishment of latency is thought to depend on complex interactions between the viral genome, the host neuronal environment, and immune surveillance mechanisms (17).

Reactivation from latency can be triggered by physiological or environmental stressors leading to renewed viral replication and spread, either peripherally or towards the central nervous system (CNS) (18). While clinically apparent reactivation typically manifests peripherally (19), increasing evidence suggests that alphaherpesviruses may also establish latency within the CNS (20, 21). Such CNS latency may result in primary or recurrent encephalitis, or in subclinical reactivation events (22).

HSV-1 is the leading cause of herpes simplex encephalitis (HSE), a severe and often fatal condition with a predilection for mesiotemporal brain regions (23, 24). Beyond encephalitis, HSV-1 has been implicated as a potential co-factor in the pathogenesis of neurodegenerative disorders such as Alzheimer’s disease (AD) (25, 26).

Although HSV-1 remains the most clinically relevant alphaherpesvirus in humans, PrV provides a valuable experimental model due to its close genetic and pathogenic relationship to HSV-1 (27, 28). In particular the attenuated mutant PrV-ΔUL21/US3Δkin, which lacks the functional tegument protein pUL21 and express a kinase-deficient pUS3, induces non-lethal encephalitis in female CD1 mice after intranasal inoculation (29). Infected animals develop severe lymphohistiocytic meningoencephalitis in mesiotemporal regions and neurological deficits reminiscent of HSE, yet most survive the acute phase, enabling long-term observation (30).

Our previous long-term studies demonstrated that infection with PrV-ΔUL21/US3Δkin results in a multiphasic disease course, with histopathological evidence of neuroinflammation, including mild meningoencephalitis and gliosis, detectable up to 168 days post-infection (dpi) (30). These findings raised key questions regarding the underlying mechanisms: whether they reflect intermittent reactivation, persistent low-level infection, chronic inflammation, or even autoimmune responses, as suggested by previous murine models and human case reports (31–33).

To address these questions, we conducted a comprehensive long-term study extending to 105 days pi. Using molecular (RNAscope™ in situ hybridization and quantitative real-time PCR (RT-qPCR)) and histopathological approaches, we systematically assessed viral gene expression, immune cell infiltration, and lesion development across defined brain regions. Given the limited number of in vivo models for PrV latency (34), we specifically evaluated the potential for CNS latency beyond the TG, with a focus on mesiotemporal structures as candidate sites of long-term viral activity.

This study integrates spatially resolved molecular and histopathological analyses to elucidate mechanisms of alphaherpesvirus persistence in the CNS and its contribution to long-term neuropathology.

## Results

The aim of this study was to identify brain regions that serve as sites for alphaherpesvirus latency and to investigate the temporal dynamics of latency and reactivation during long-term infection.

Female CD1 mice (6-8 weeks old) were intranasally inoculated and sacrificed at 21, 42 and 105 dpi for RT-qPCR, RNA in situ hybridization, and histopathological analyses (study 1). Archived brain sections from a previous long-term study (study 2) using the same model (30) were reanalyzed to investigate the post-acute phase at 28 dpi. In that study, a subset of animals received cyclophosphamide/dexamethasone at 170 dpi and were sacrificed 20 days later. These tissues were included for RNA in situ hybridization and histopathology. An overview of the experimental design is shown in Fig 1.

**Fig 1.**
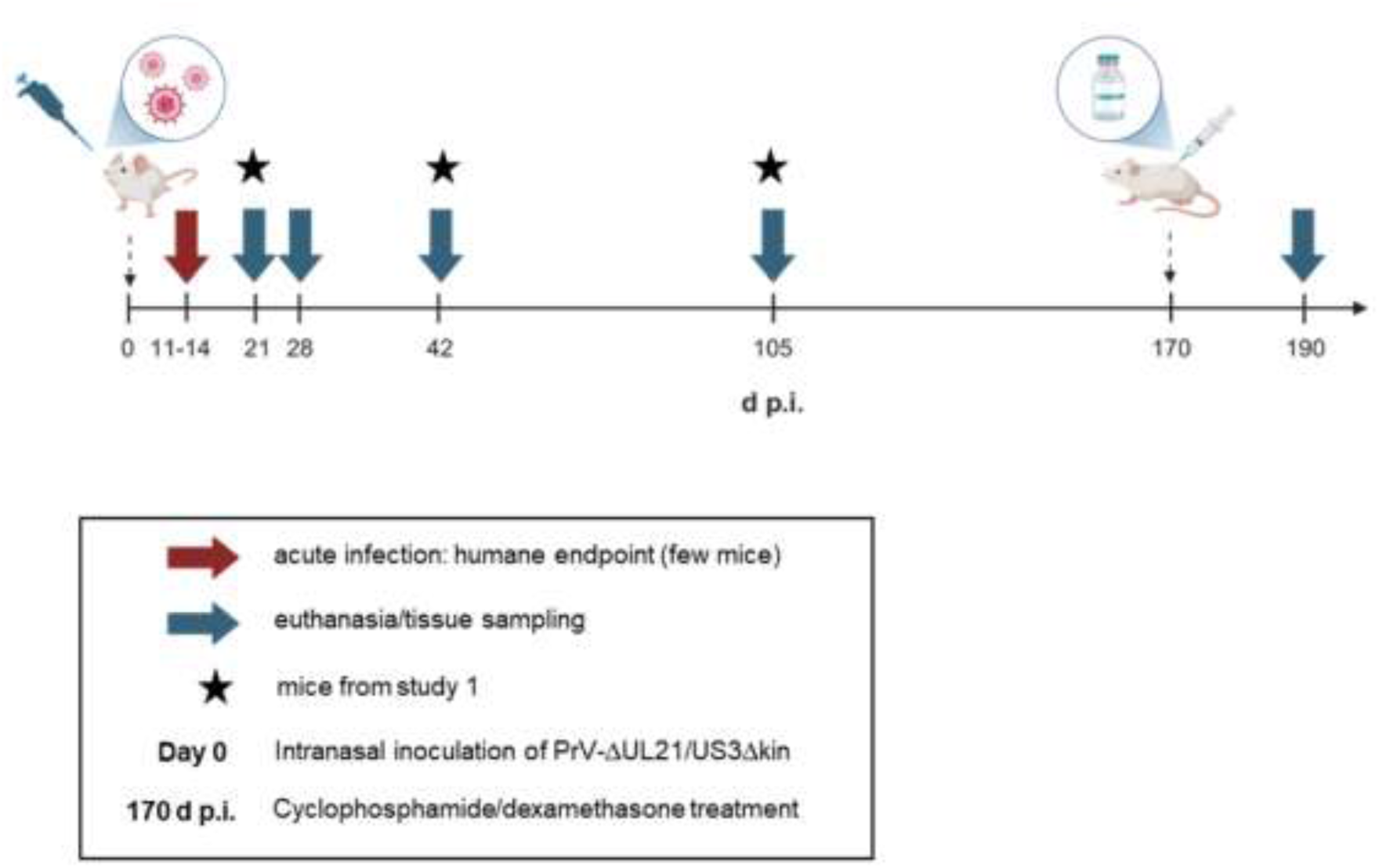
Experimental timeline of necropsies and tissue collection. Six-to eight-week-old CD1 mice were intranasally inoculated with PrV-ΔUL21/US3Δkin (day 0) in two independent studies. In study 1, tissue samples were collected at 21, 42, and 105 dpi (indicated by stars). In study 2, mice received an immunosuppressive treatment (cyclophosphamide/dexamethasone) at 170 dpi, and tissue was collected at 28 dpi and 190 dpi. During the acute infection phase (11–14 dpi), mice that reached the humane endpoint were euthanized, and tissues were collected for analysis. Created with BioRender.com

### Clinical evaluation

As previously reported in mice from study 2 (0-168 dpi) (30), infection with PrV-ΔUL21/US3Δkin in study 1 followed a multiphasic disease course. From day 5 post-infection, mice developed mild clinical signs including ruffled fur and reduced activity. Disease symptoms peaked between 10-15 dpi, with incidence rates approaching 90%. During this acute phase, mice displayed alopecic skin erosions, nasal bridge edema, mild pruritus, and behavioral abnormalities such as hyperactivity and “star gazing”. Approximately 10% of animals reached humane endpoint criteria due to severe manifestations including seizures and hyperexcitability.

By 30 dpi, predominantly mild clinical signs persisted in ∼50% of mice. Between 45 and 105 dpi, incidence declined further to 20–30%, with fluctuating low-grade signs such as ruffled fur and subtle behavioral alterations (Fig 2A).

**Fig 2.**
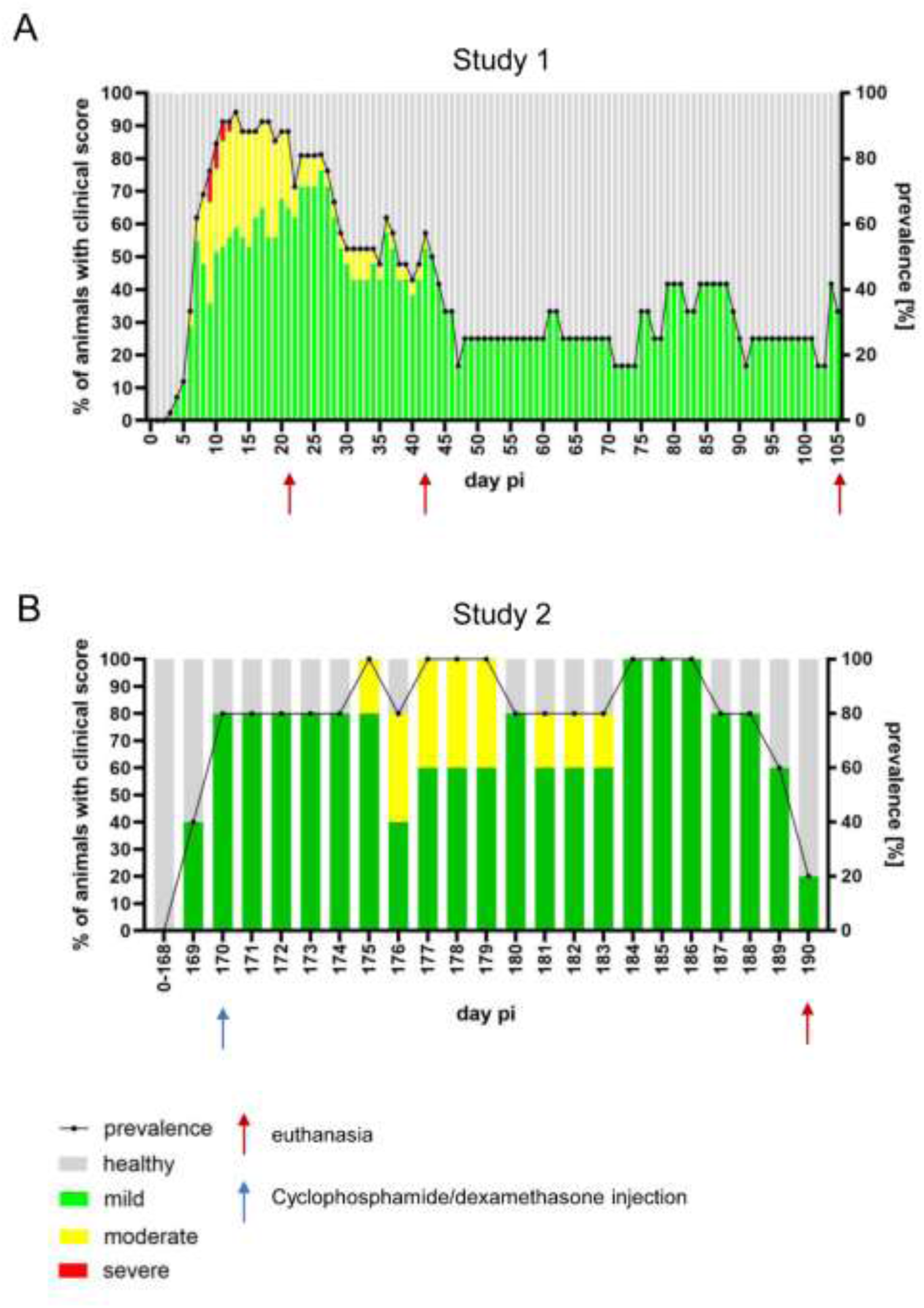
Longitudinal assessment of disease severity and incidence in PrV-ΔUL21/US3Δkin-infected mice. The proportion of clinically affected animals is shown as a percentage of the total number of infected mice. Disease severity is color-coded as follows: healthy (grey), mildly affected (green), moderately affected (yellow), and severely affected (red). Percentages are plotted on the left y-axis, while the overall incidence of clinical signs is indicated by a dotted line and shown on the right y-axis. Red arrows indicate necropsy time points. (A) Study 1: PrV-ΔUL21/US3Δkin or mock-inoculated mice (n=7) were sacrificed at 21, 42, and 105 dpi. (B) Study 2: Mice (n=5) were inoculated, monitored, and evaluated as in study 1, but received immunosuppressive treatment with cyclophosphamide/dexamethasone at 170 dpi (blue arrow), and were euthanized at 190 dpi.

Following cyclophosphamide/dexamethasone treatment at 170 dpi in study 2, approx. 80% of animals developed mild clinical signs (ruffled fur, reduced activity) within the first 5 days post treatment (dpt). Between 5 and 10 dpt, the disease severity increased, and all animals exhibited signs of reactivation. Approximately 50% developed moderate symptoms including “star gazing,” pruritus, and mild hunching. By 20 dpt, both incidence and severity steadily declined, with approx. 10% of animals were still symptomatic at the end of the observation period (Fig 2B).

### Expression patterns of lytic and latency-associated transcripts

To characterize spatial and temporal expression of viral transcripts, brains from mice sacrificed at 11– 14 dpi (reached humane endpoint), 28, 42, 105, and 190 dpi were analyzed. In situ RNA hybridization was performed on six standardized coronal brain sections per animal (Fig 3A, L1–L6), using the RNAscope™ method (35) with probes targeting the lytic gene UL19 and the latency-associated transcript LAT (Fig 3B).

**Fig 3.**
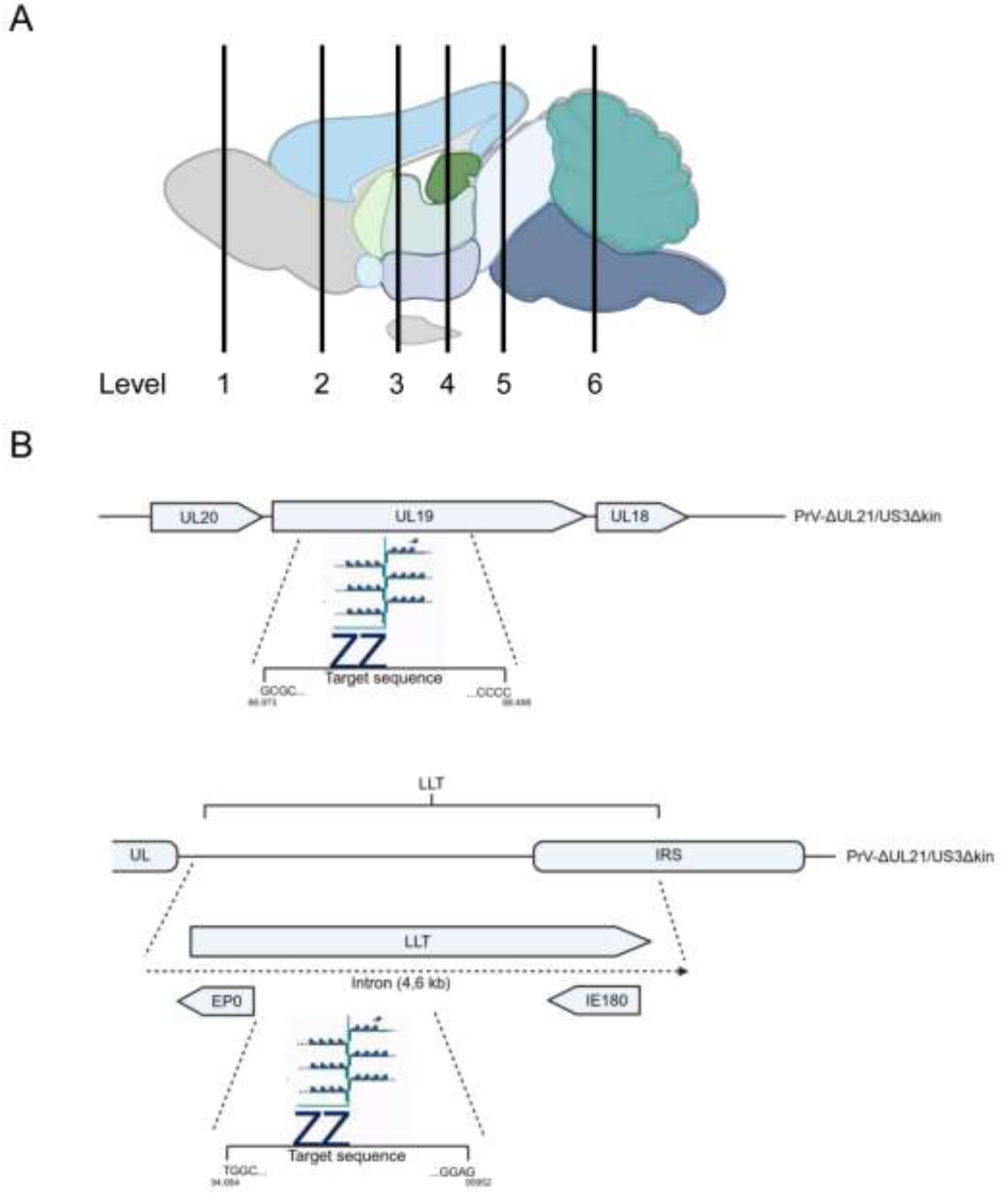
Experimental setup for RNAscope™ analysis. (A) Schematic overview of analyzed brain regions. Serial 3 µm coronal sections were collected for detailed analysis at six defined anatomical levels: L1 (olfactory bulb), L2 (prefrontal cortex), L3 (frontoparietal cortex, basal ganglia), L4 (parietal cortex, thalamus, hypothalamus, hippocampus), L5 (midbrain), and L6 (cerebellum/pons). (B) Design of UL19 (V-SHSV-UL19-C1) and LLT (V-SuHV1-LLT-O2-C2) probes used for detection of lytic and latency-associated (LAT) viral mRNA. V-SHSV-UL19-C1 targets the UL19 and V-SuHV1-LLT-O2-C2 the LAT region, which is represented by the blue dots and connecting lines. Location of the probes corresponds to nucleotide positions: 66973–68498 base pairs (UL19), 94664-95952 base pairs (LLT). Created with BioRender.com.

#### CNS sites of LAT expression

LAT signal was detected on multiple CNS structures across time points (Fig 4). In the hindbrain, expression was observed in the cerebellum (Cb) and brainstem (BS) nuclei including the nucleus of the solitary tract (Sol), inferior olive (IO), spinal trigeminal nucleus (Sp5), spinal vestibular nucleus (SpVe), facial nucleus (7N), and the reticular formation (FR).

**Fig 4.**
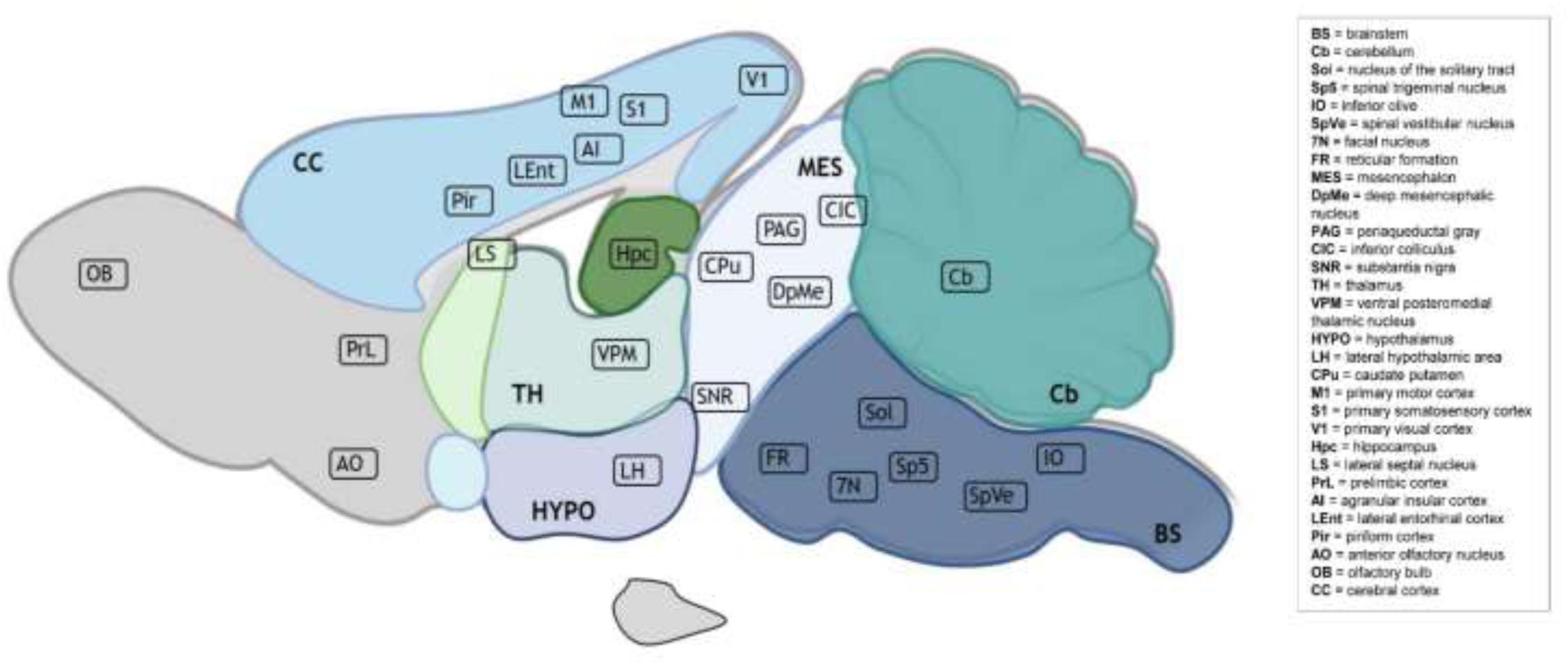
Widespread detection of latency-associated transcript (LAT) expression in the murine brain following intranasal PrV-ΔUL21gfp/US3Δkin infection. LAT RNA was detected by RNAscope™ in all analyzed anatomical regions across functional brain systems (brainstem, mesencephalon, diencephalon, telencephalon, and cerebellum) at 11–14, 28, 42, 105, and 190 dpi. The distribution of LAT-positive regions are illustrated on a schematic sagittal brain section. Created with BioRender.com.

In the midbrain, LAT-positive signals were found in the substantia nigra (SNR), caudate putamen (CPu), deep mesencephalic nucleus (DpMe), and central nucleus of the inferior colliculus (CIC). Thalamic and hypothalamic areas such as the ventral posteromedial nucleus (VPM) and lateral hypothalamus (LH) showed expression.

In the telencephalon, signals were consistently detected in mesiotemporal regions including the hippocampus (Hpc), lateral septal nucleus (LS), and lateral entorhinal cortex (LEnt). Additional expression was observed in the piriform cortex (Pir), agranular insular cortex (AI), prelimbic cortex (PrL), anterior olfactory nucleus (AO)), as well as in primary sensory (S1), visual (V1), and motor cortices (M1), and in the olfactory bulb (OB).

#### Semi-quantitative analysis of UL19 and LAT expression

To assess expression dynamics, nine brain regions were selected for semi-quantitative analysis at defined post-infection timepoints. Selected regions included known alphaherpesvirus target areas such as Pir, AI, LEnt, Hpc, S1, Sp5 and OB (36, 29). The Cb served as a reference region due to its low susceptibility to alphaherpesviral infection (37). The TG was included as a canonical latency site (1) (Fig 5A).

**Fig 5.**
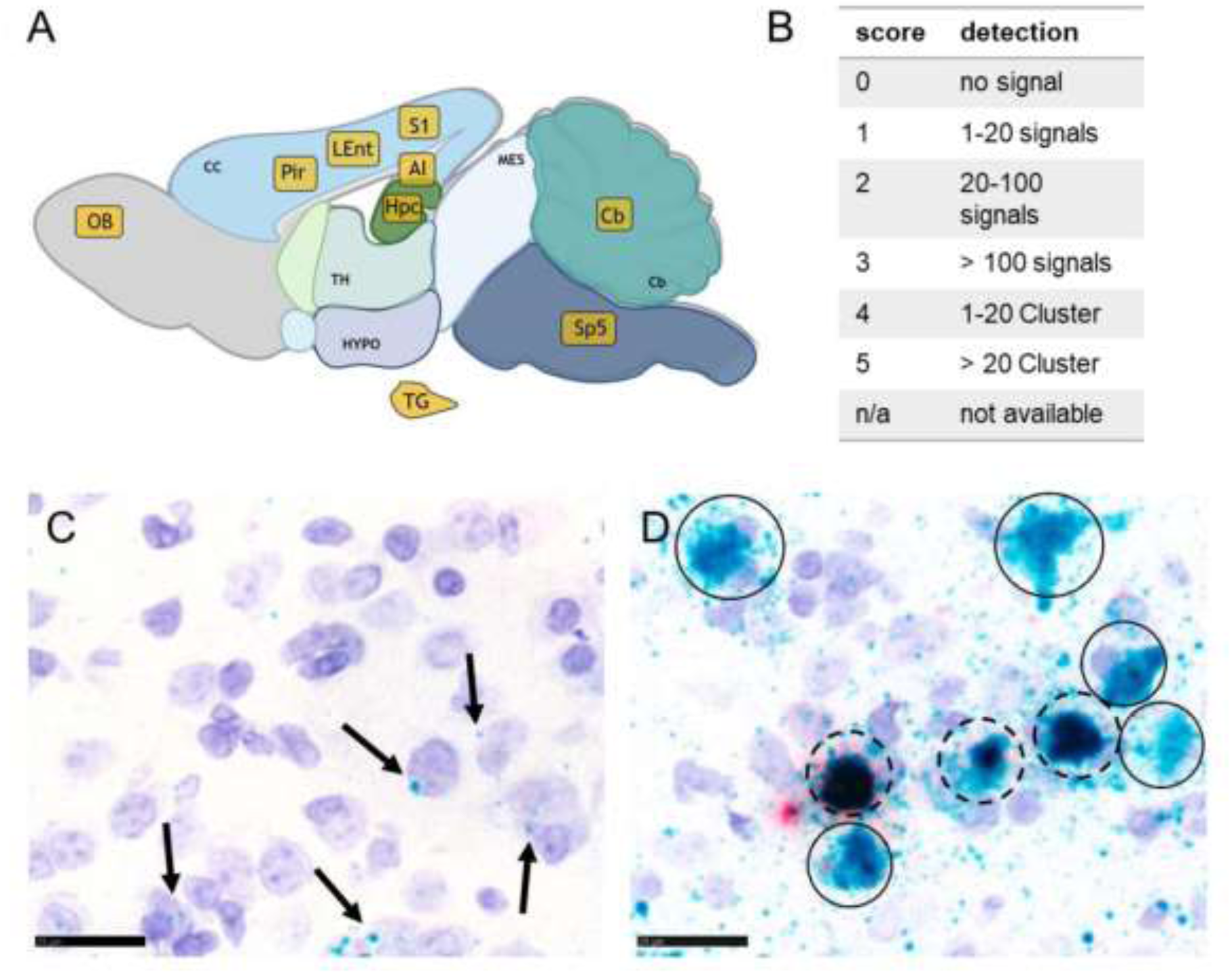
Detection of UL19 and LAT RNA transcripts in murine brain tissue using RNAScope™ in situ hybridization. (A) Overview of the selected brain regions analyzed (yellow-shaded boxes). (B) Scoring criteria for positive signal detection based on the number and clustering of signals within these regions. Coronal brain sections representing six anatomical levels were examined by light microscopy. If multiple levels were available, the highest score per anatomical region was recorded. (C) Individual RNA transcripts appear as distinct chromogenic dots (arrow); green dots represent UL19 RNA transcripts. (D) Accumulation of transcripts may result in clusters (solid circles), which can comprise different transcripts, indicated by a mixed red (LAT) and green (UL19) signals (dashed circles). TG = trigeminal ganglion, Cb = cerebellum, Sp5 = spinal trigeminal nucleus, S1 = primary somatosensory cortex, Hpc = hippocampus, LEnt = lateral entorhinal cortex, Pir = piriform cortex, AI = agranular insular cortex, OB = olfactory bulb. Scale bar = 25 µm. Created with BioRender.com.

Signals were scored on a 0-5 scale (Fig 5B), distinguishing discrete puncta (Fig 5C) from clustered signal patterns (Fig 5D). Results are summarized in Fig 6 and detailed per animal in S1 Table. Representative RNAscope™ images are shown in Fig 7 and Fig 8.

**Fig 6.**
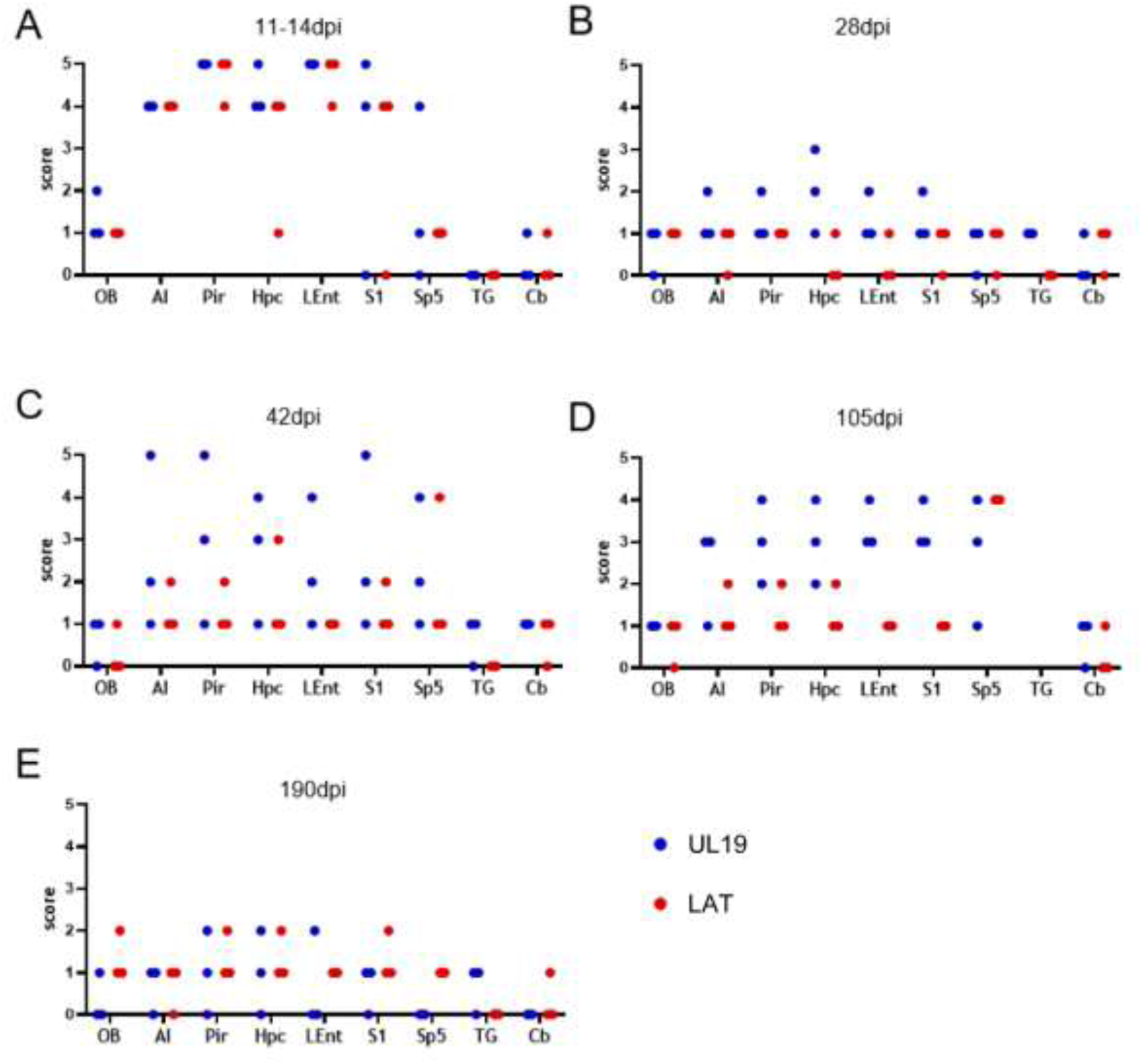
Semi quantitative analysis of RNAScope™ signal detection in PrV-infected murine brain tissue. Positive signals for UL19 (blue) and LAT (red) RNA transcripts were scored at 11-14 dpi (A), 28 dpi (B), 42 dpi (C), 105 dpi (D) and 190 dpi (E). Scores represent data from three mice per time point and brain region. TG = trigeminal ganglion, Cb = cerebellum, Sp5 = spinal trigeminal nucleus, S1 = primary somatosensory cortex, Hpc = hippocampus, LEnt = lateral entorhinal cortex, Pir = piriform cortex, AI = agranular insular cortex, OB = olfactory bulb.

**Fig 7.**
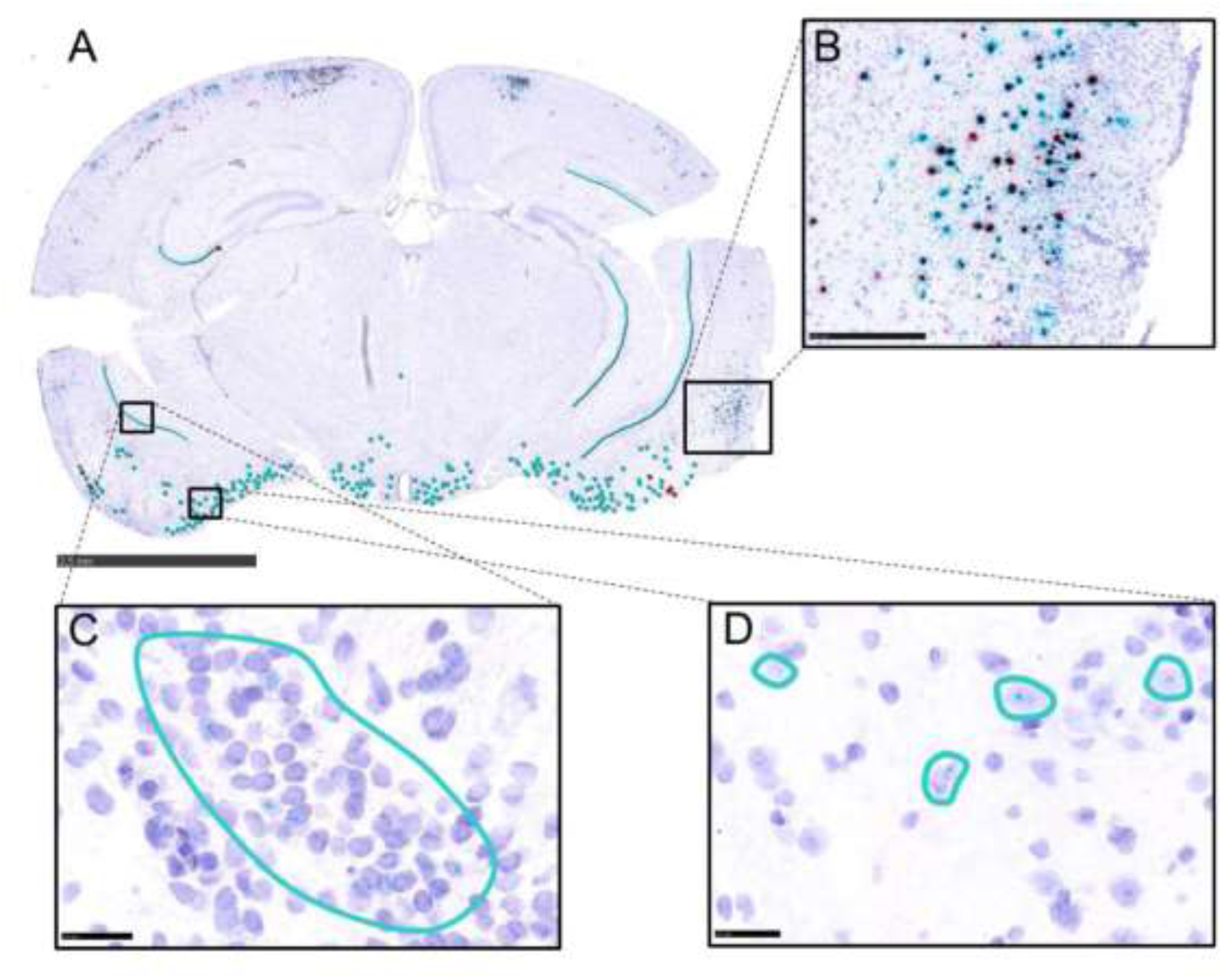
Detection of UL19 and LAT RNA transcripts during the acute phase of PrV-ΔUL21/US3Δkin infection using RNAscope™ in situ hybridization. (A) Coronal section of a murine brain at the level of the temporal lobe showing UL19 (green dots) and LLT (red dots) RNA signals at 14 dpi with PrV-ΔUL21/US3Δkin. Scale bar = 2,5mm (B) Higher magnification of UL19 and LAT signals in LEnt. Scale bar = 250 µm. (C) Higher magnification of scattered UL19 signals in Hpc (cyan outline). Scale bar = 25µm. (D) Higher magnification of UL19 signals in Pir (cyan circles). Scale bar = 25µm. Hpc = hippocampus, LEnt = lateral entorhinal cortex, Pir = piriform cortex.

**Fig 8.**
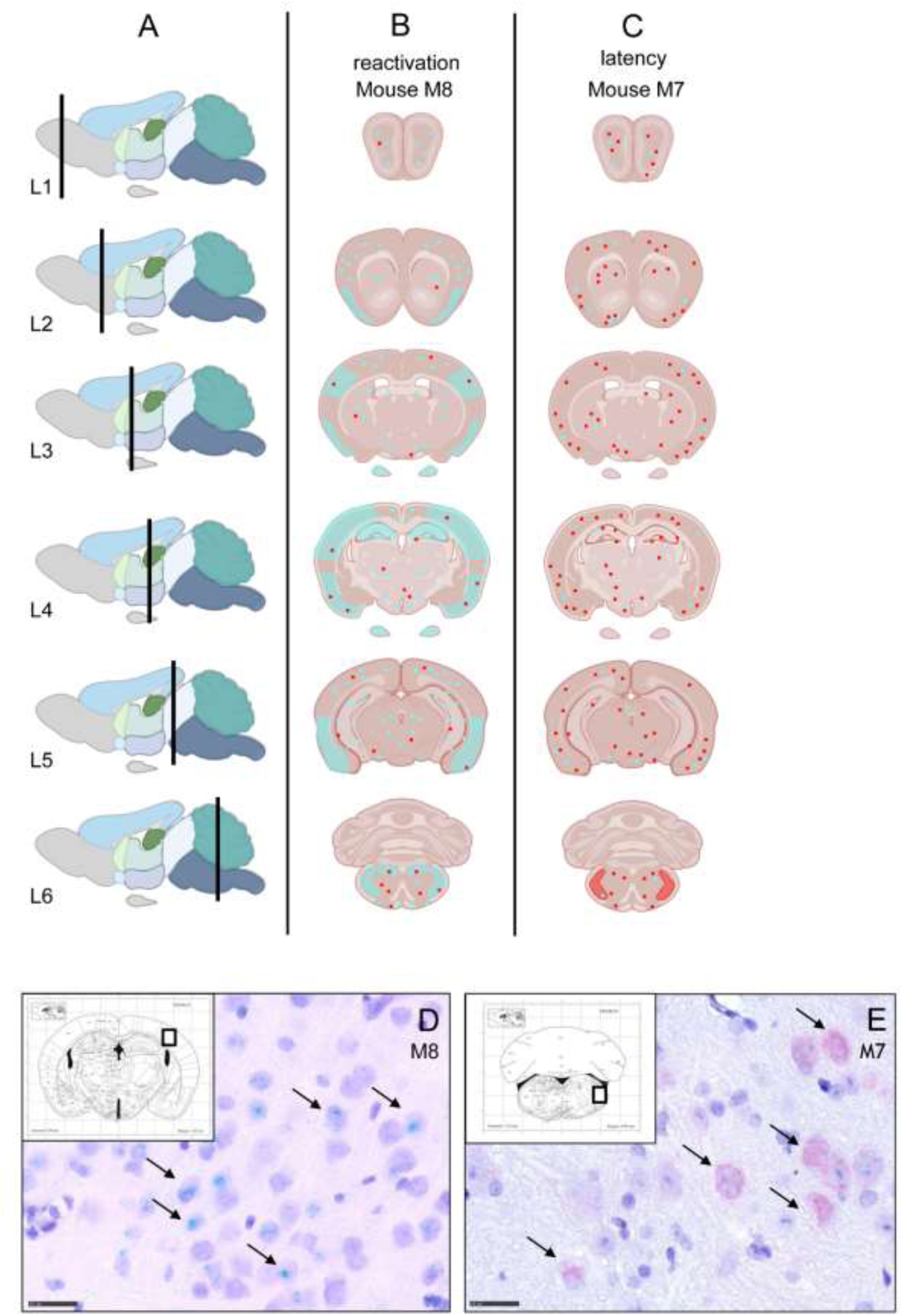
Distribution patterns of UL19 and LAT RNA signals across defined anatomical brain regions in selected mice at 42 dpi. (A) Sagittal view of the murine brain indicating the coronal section levels analyzed (L1-6). (B) Schematic coronal sections of mouse M8 during viral reactivation, depicting UL19 and LAT distribution. (C) Schematic coronal sections of mouse M7 during latency, depicting UL19 and LAT distribution. In panels B and C, dots indicate single RNA transcripts, while shaded areas represent clusters (red = LAT, cyan = UL19). (D) Higher magnification of the primary somatosensory cortex (S1) in mouse M8 showing intense intraneuronal UL19 signals, with clusters indicated by arrows. (E) Higher magnification of the spinal trigeminal nucleus (Sp5) in mouse M7, showing intense neuronal LAT signals, with clusters indicated by arrows. Sp5 = spinal trigeminal nucleus, S1 = primary somatosensory cortex. Scale bar = 25 µm. Coronal brain sections shown with *The Mouse Brain in Stereotaxic Coordinates* by Paxinos and Franklin, 2001. Created with BioRender.com.

11-14 dpi:

Both, UL19 and LAT transcripts were highly expressed (Fig 6A), particularly in temporal and frontoparietal regions (Fig 7A). Strong signals (Score 4–5) were observed in the LEnt; (Fig 7B), Hpc (Fig. 7C), Pir (Fig. 7D), AI, and S1. Sp5 and OB showed only few signals (Score 1). In the Cb only minimal or no signals (Score 0–1) were present, and no transcripts were detected in the TG (Fig 6A).

28 dpi:

LAT expression was at baseline across all regions (Score 0-1), with no signal in the TG. UL19 expression remained low in OB, Sp5, TG, and Cb, but was modestly elevated in the LEnt, Pir, AI, S1, and Hpc (Score 1-2) (Fig 6B).

42 dpi:

LAT largely remained at baseline, except in one animal showing increased signals (AI, Pir, S1: score 2; Hpc: score 3; Sp5: score 4). UL19 expression was very heterogenous. Cb, TG, and OB remained low (score 0–1), whereas AI, Pir, Hpc, LEnt, S1 and Sp5 varied widely (1–5) (Fig 6C). Representative animals illustrate this: Mouse M8 displayed UL19 clusters across regions, consistent with reactivation (Fig 8B), while mouse M7 showed LAT clusters in Sp5 and increased signal frequency in AI, Pir, Hpc, and S1, with low UL19 expression, suggestive for latency (Fig 8C). Representative RNAscope™ images of clustered transcript signals in M8 (S1) and M7 (Sp5) are provided in Fig 8D and Fig 8E, respectively.

105 dpi:

LAT remained at baseline, except consistently elevated signals in Sp5 (Score 4). UL19 again showed heterogeneity, with OB and Cb low (score 0 and 1), and highest variability in Sp5 (score 1–4), AI (1–3), Pir/Hpc (2–4), and LEnt/S1 (3–4) (Fig 6D). No TG samples were available for this time point.

190 dpi (post-immunosuppression):

LAT remained mostly at basal levels with isolated increases in S1, Hpc, Pir, and OB (score up to 2). UL19 was generally low (score 0-1), with isolated signals in Pir, Hpc, and Lent (up to score 2). (Fig 6E).

A shift in the intracellular distribution of viral transcripts was observed. During the acute phase, UL19 and LAT were frequently detected within the same neuron or in immediately adjacent neurons. From 28 dpi onward and persisting through the end of the observation period, this pattern changed and both UL19 and LAT were detected in different cells with no co-localization in defined neurons.

### Quantification of viral DNA by RT-qPCR

To complement RNAscope™ findings, RT-qPCR was performed on six brain regions (OB, Pir, temporal lobe (TL), TG, Cb, BS) from mice sacrificed at 9-10 (humane endpoint), 21, 42 and 105dpi. Ct values for UL19 and LAT are shown in Fig 9. Clinical correlations per mouse are detailed in S1 Fig.

**Fig 9.**
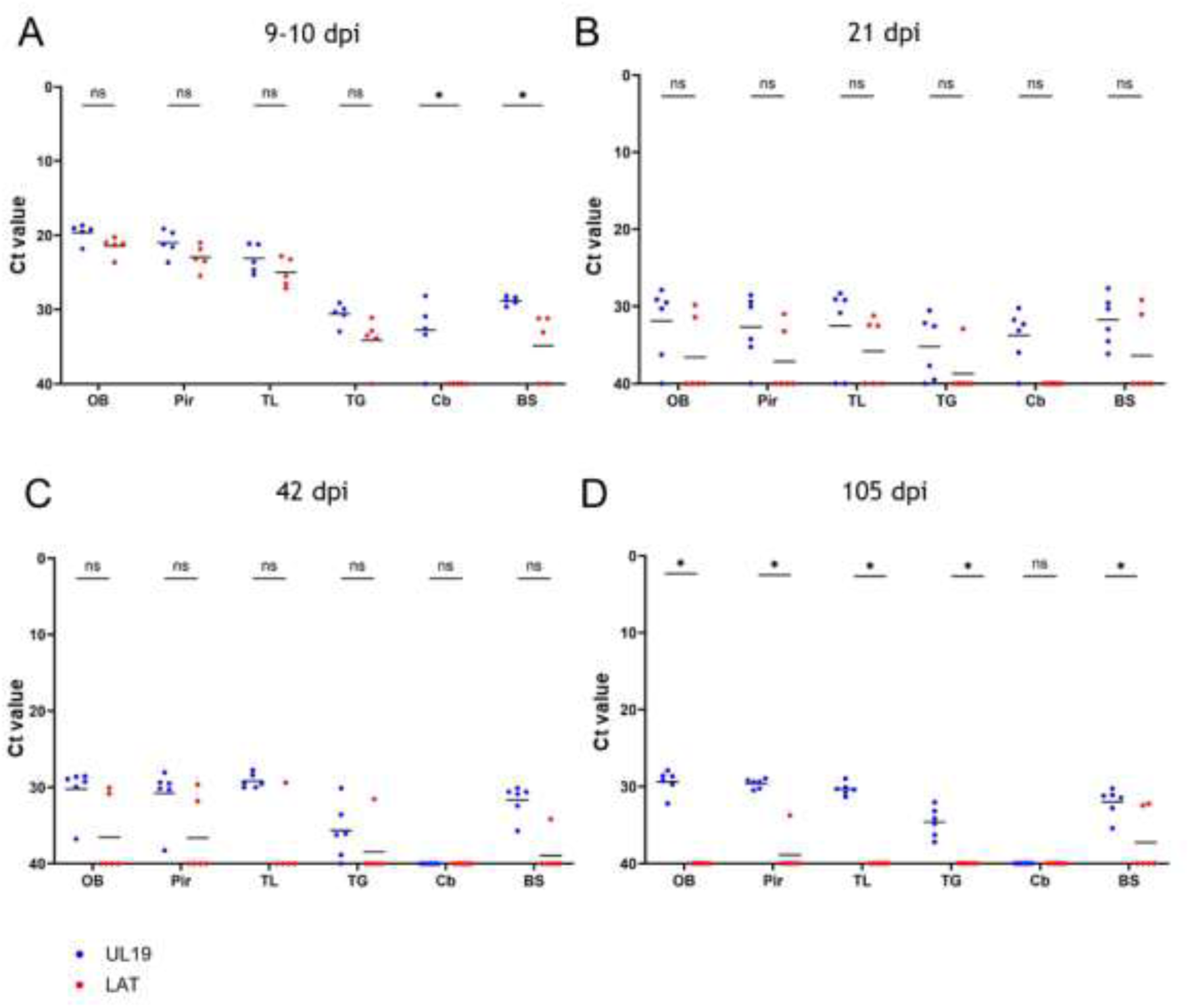
Quantification of UL19 and LAT genomic DNA in selected brain regions using RT-qPCR. cDNA levels of the lytic gene UL19 and the latency-associated transcript (LAT) were quantified by RT-qPCR in brain tissue from six mice per time point. Tissue was collected at four time points: (A) 9-10 dpi (five animals euthanized at the humane endpoint), (B) 21 dpi, (C) 42 dpi, and (D) 105 dpi. Dissected brain regions included the olfactory bulb (OB), piriform cortex (Pir), temporal lobe (TL), trigeminal ganglion (TG), cerebellum (Cb) and brainstem (BS). Ct values are shown for each region. Bars represent the geometric mean per group. *p<0,001.

9-10 dpi:

Animals reaching the humane endpoint exhibited severe signs (hunching, seizures, >20% weight loss) (S1 Fig). High expression levels of both UL19 and LAT were detected in the OB, TL, and Pir, as indicated by low Ct values, reaching values below 25. In contrast, the TG, Cb, and BS showed higher Ct (> 30), indicating lower abundance (Fig 9A).

21 dpi:

Both transcripts were near the detection threshold (Ct > 35). UL19 signals tended to be slightly more abundant than LAT, though not significant. Two mice (M25 and M24) showed lower LAT Ct values; but only one (M25) displayed seizures and localized alopecia (Fig. 9B, S1 Fig).

42 dpi:

UL19 expression was modestly elevated in OB, TL, Pir, and BS (Ct > 30), but undetectable in Cb and TG. LAT was absent except in one mouse (M29), which displayed low-level expression in all regions except Cb (Fig 9C).

105 dpi:

UL19 remained detectable in all regions except Cb (Ct 28-35). LAT was largely absent, except in two animals (M36, M37) with low-level expression in BS and Pir (Ct 32) (Fig 9D). Both showed only mild clinical signs (ruffled fur) (S1 Fig).

### Histopathological temporal profiling

Histopathology was performed on the same brain tissue analyzed by RNAscope™.

#### Long-term histomorphological alterations in the CNS

Histomorphological changes were consistently observed throughout the entire observation period across the analyzed brain regions (Fig 10).

**Fig 10.**
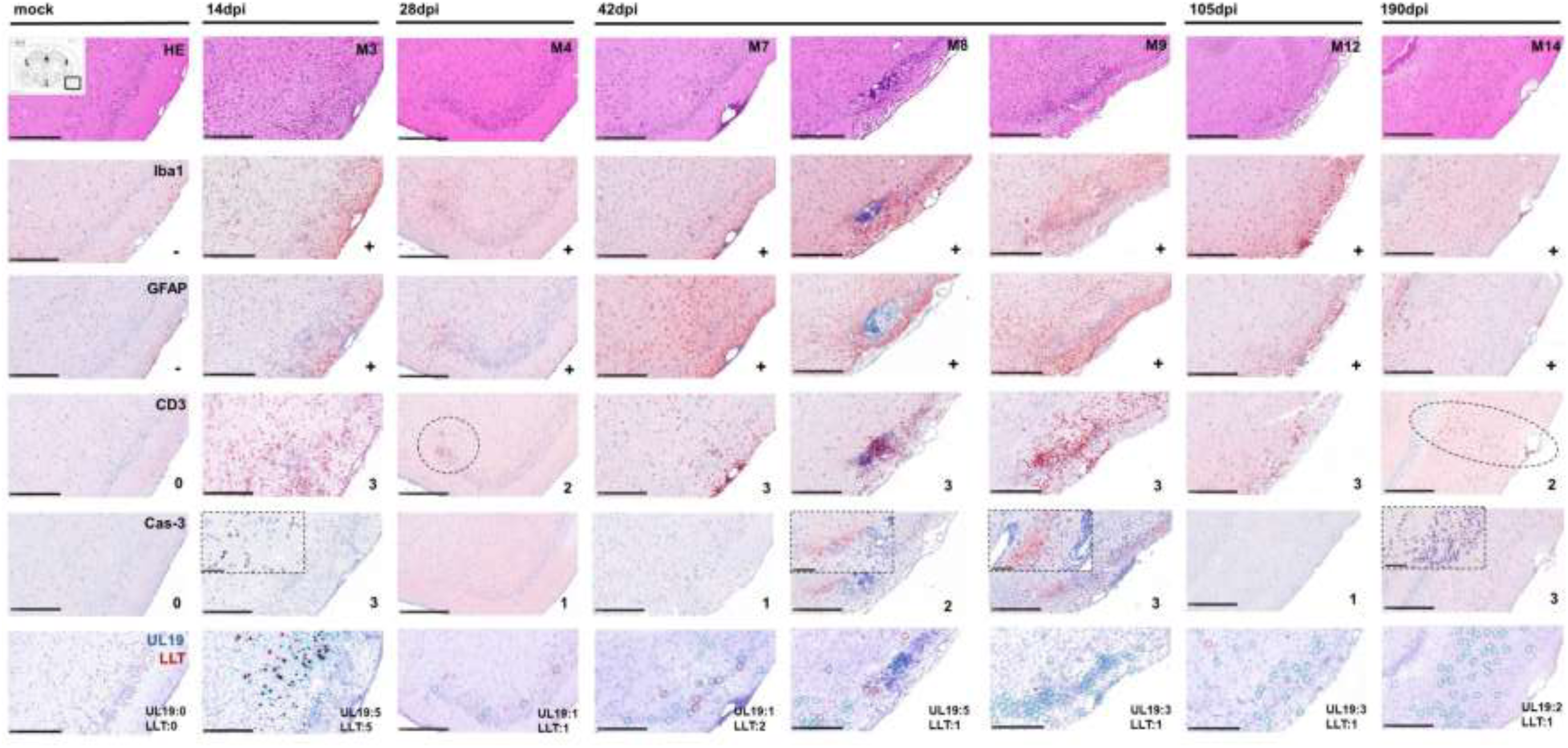
Representative images of long-term lesions in the temporal lobe (piriform cortex) from 14 to 190 dpi in PrV-ΔUL21/US3Δkin-infected mice. Temporal progression of infection-associated neuropathology was assessed by histopathology (H&E), immunohistochemistry (Iba1, GFAP, CD3, cleaved caspase-3 (Cas-3), and RNAscope™ in situ hybridization targeting UL19 (lytic) and LAT (latency-associated) transcripts. Representative brain sections are shown for mice sacrificed 14, 28, 105, and 190 dpi; three animals are shown for 42 dpi to illustrate interindividual variability histopathological and molecular findings. At 14 dpi, severe necrotizing meningoencephalitis was evident in the temporal lobe, accompanied by dense infiltrates of CD3⁺ T-cells, Iba1⁺ microglia/infiltrating macrophages, reactive GFAP⁺ astrocytes, and abundant Cas-3⁺ apoptotic cells. From 28 to 105dpi, mild meningoencephalitis was detected, characterized primarily by CD3+ and Iba1+ immune cells. A gradual increase in GFAP expression indicated ongoing astrogliosis, while only a few Cas-3⁺ cells were detected, suggesting reduced apoptotic activity. At 42dpi, mild to moderate meningoencephalitis was present, with pronounced CD3⁺ T-cell infiltration and interindividual variability in Cas-3⁺ cell density ranging from low to high. At 190dpi, mild meningoencephalitis was still evident, characterized by ongoing CD3⁺, Iba1⁺ and GFAP⁺ immune cell infiltration together with a high number of Cas-3⁺ cells. Semi-quantitative scores from IHC and in situ hybridization analyses of the piriform cortex (ROI) are shown in the lower right corner of each panel. Scale bar = 250 µm.

H&E staining revealed severe necrotizing meningoencephalitis during the acute phase, predominantly affecting mesiotemporal regions (piriform and prefrontal cortices). Lesions were associated with dense T-cell and macrophage infiltrates, neuronal necrosis, and marked gliosis. Between 28 and 190 dpi, pathology shifted to a milder phenotype. Low-grade meningoencephalitis persisted with scattered single-cell necrosis in mesiotemporal and frontal regions. Perivascular and meningeal T-cell/histiocyte infiltrates remained detectable, alongside ongoing glial activation (S1 Table).

#### Spatiotemporal T cells dynamics and correlation with lytic transcription

Region-specific patterns of T-cell infiltration detected by CD3 staining were observed (Fig 11A). In Pir and LEnt, T-cell scores remained high (2–3) through 105 dpi, declining slightly (1–2) at 190 dpi. AI showed a similar trajectory with earlier decline. Hpc infiltration remained at score 2–3 at most time points, peaking at 190 dpi. S1 fluctuated, with high scores at 11–14, 42, and 105 dpi. Sp5 was largely unaffected, except for a transient peak at 11–14 dpi. TG, Cb, and OB consistently showed minimal infiltration. Correlation analysis demonstrated a strong positive association between CD3⁺ T-cell density and UL19 expression, particularly at 11–14 dpi (p ≤ 0.05) and 42 dpi (p ≤ 0.01) (Fig 11B).

**Fig 11.**
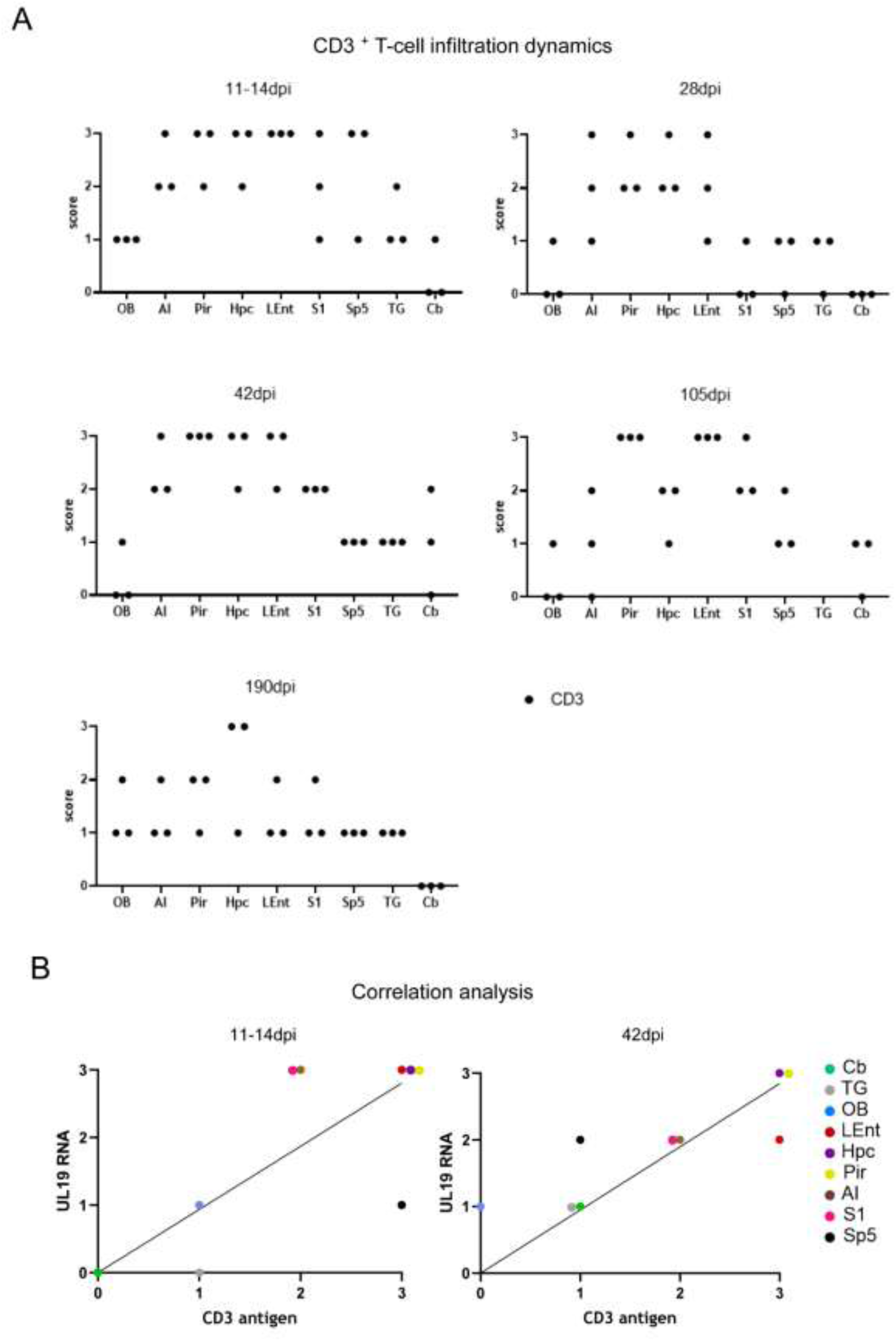
Semiquantitative analysis of CD3⁺ T-cell infiltration and correlation with UL19 RNA in PrV-ΔUL21/US3Δkin-infected murine brain tissue. (A) Semiquantitative scoring of CD3^+^ T-cell infiltration in selected brain regions at 11-14, 28, 42, 105 and 190 dpi. Scores from three mice per time point are shown. (B) Correlation analysis between CD3⁺ T-cell infiltration and UL19 RNA transcript detection. Mean CD3 antigen scores (x-axis) and mean UL19 RNA scores (y-axis) from three replicates per region were subjected to linear regression and Spearman’s correlation analysis. A significant positive correlation was observed at 11–14 dpi (p ≤ 0.05) and 42 dpi (p ≤ 0.01). TG = trigeminal ganglion, Cb = cerebellum, Sp5 = spinal trigeminal nucleus, S1 = primary somatosensory cortex, Hpc = hippocampus, LEnt = lateral entorhinal cortex, Pir = piriform cortex, AI = agranular insular cortex, OB = olfactory bulb.

### Integrated spatiotemporal profiling of viral transcripts, neuroinflammation, and clinical outcome

Representative images from the piriform cortex are shown in Fig 10.

During the acute phase (11–14 dpi), animals developed severe meningoencephalitis characterized by dense T-cell infiltrates (score 3), marked gliosis, and abundant apoptotic cells (cleaved caspase-3 (Cas-3) score 3). Strong signals for both UL19 and LAT transcripts (score 5) were detected, coinciding with severe clinical manifestations.

At 28 dpi, inflammation had subsided to a mild meningoencephalitis, accompanied by ongoing gliosis. T-cell infiltration (score 2) and apoptotic activity (Cas-3 score 1) were reduced, and viral transcript levels were low (score 1). Correspondingly, only minimal clinical signs were observed.

At 42 dpi, interindividual variability became apparent. In mouse M7, UL19 expression was low (score 1), LAT moderately increased (score 2), and apoptotic activity minimal. In contrast, M8 exhibited strong UL19 expression (score 5), moderate apoptosis (Cas-3 score 2), and low LAT, consistent with viral reactivation. Mouse M9 showed moderate UL19 levels (score 3), pronounced apoptosis (Cas-3 score 3), and behavioral abnormalities such as stargazing.

At 105 dpi, mild meningoencephalitis persisted, with high T-cell infiltration (score 3), moderate UL19 expression (score 3), and low LAT (score 1). Apoptosis remained minimal (Cas-3 score 1), while overt clinical signs were still present.

By 190 dpi, inflammation was mild, with ongoing T-cell infiltration (score 2) and low expression of UL19 (score 2) and LAT (score 1). Notably, apoptotic activity was elevated (Cas-3 score 3), despite the absence of clinical signs.

Mock-infected mice showed no abnormalities. An immunosuppressed mock control, however, exhibited increased apoptosis (Cas-3 score 3). A full summary of findings is provided in S1 Table.

## Discussion

In this study, we examined the long-term dynamics of alphaherpesvirus infection in the murine CNS after intranasal inoculation with PrV-ΔUL21/US3Δkin. By integrating molecular, histopathological, and clinical analyses, we identified distinct spatiotemporal patterns of lytic and latent viral transcription and linked them to neuroinflammation and clinical outcome. In addition, an immunosuppressed cohort allowed us to assess the potential for viral reactivation.

The infected mice followed a multiphasic disease trajectory consistent with previous observations (30). During the acute phase, severe clinical signs including seizures, hyperactivity, and “stargazing” were observed, reflecting neuronal dysfunction particularly in mesiotemporal regions (38, 39). Mild to moderate signs persisted in a subset of animals during later time points, and immunosuppression at 170 dpi induced transient disease recurrence. These manifestations closely resemble the clinical signs reported in human herpes simplex encephalitis (HSE), where hippocampal and entorhinal involvement underlies cognitive deficits, memory impairment, and seizures (22, 40, 41).

RNAscope™ analysis demonstrated widespread neuronal permissiveness to latency, with LAT signals present across hindbrain, midbrain, diencephalon, and telencephalon including mesiotemporal, olfactory, and neocortical regions. Semi-quantitative analysis revealed distinct temporal patterns: during the acute phase (11–14 dpi), both LAT and UL19 were abundant in mesiotemporal and frontoparietal areas, by 28 dpi both signals had declined; and from 42 dpi onward, LAT remained at baseline, whereas UL19 displayed interindividual heterogeneity, with some animals showing strong signals suggestive of reactivation. Following immunosuppression (190 dpi), LAT remained low, while UL19 expression showed modest increases in selected regions.

RT-qPCR confirmed these dynamics, showing highest viral loads in the olfactory bulb, temporal lobe, and piriform cortex during the acute phase, with lower but persistent UL19 detection at later stages. Discrepancies between RNAscope™ and RT-qPCR, particularly in the OB, likely reflect methodological differences: whole-tissue qPCR versus single-section histological analysis.

A notable finding was the absence of detectable LAT signals in the TG, despite its canonical role as the primary latency reservoir (1, 42, 10). Instead, brainstem regions, especially the Sp5, consistently exhibited LAT expression at later time points. This aligns with reports suggesting that reactivation may occur more readily in brainstem neurons than in the TG (43, 20).

During the acute phase, LAT and UL19 were often detected within the same neuron, suggesting transient overlap of lytic and latent transcription. From 28 dpi onward, however, transcripts were strictly segregated to distinct cells, with no further co-localization. This shift suggests an early overshooting response that subsequently resolves into mutually exclusive transcriptional programs at the single-cell level. A similar phenomenon has also been described by Zhang ((44)), and is consistent with evidence that lytic and latent gene expression can transiently coexist (45–48).

UL19 expression showed pronounced inter-individual variability at 42 and 105 dpi. Some animals displayed strong UL19 signals across multiple regions, consistent with reactivation paralleling findings from Menendez ((49)), where lytic HSV-1 activity was detected post-60 dpi, while others exhibited primarily LAT signals with sparse UL19 expression, suggestive of latent infection. Such variability may reflect differences in latent genome copy numbers, which correlate with reactivation frequency (50, 51).

Histopathological analysis confirmed persistent CNS inflammation. During the acute phase, severe necrotizing meningoencephalitis was observed in mesiotemporal, piriform, and prefrontal regions. Thereafter, mild but sustained lymphohistiocytic meningoencephalitis, gliosis, and focal neuronal necrosis was present up to end of the experiment (190 dpi). Strong T-cell infiltration followed region-specific patterns and correlated with UL19 expression, particularly at 11–14 and 42 dpi, supporting the concept that T-cell clusters mark localized reactivation sites (42). CD8⁺ T cells are known to suppress reactivation in an antigen-specific manner (47), and LAT transcripts may counteract this by inhibiting apoptosis through caspase-3 regulation (52). In our study, cleaved caspase-3 was frequently detected in regions with high UL19 expression and strong T-cell infiltration, further supporting the link between lytic activity, immune surveillance, and apoptosis.

Together these findings support two non-exclusive interpretations: (i) episodic, often subclinical reactivation, reflected by interindividual peaks of UL19 expression and clustered T-cell infiltrates, and (ii) chronic low-grade infection, indicated by persistent inflammation and ongoing UL19 detection across most animals and time points. This duality is consistent with reports of both spontaneous reactivation and chronic neuroinflammation during HSV-1 latency (20, 53, 31, 32, 54).

Beyond direct viral activity, secondary mechanisms may also contribute to long-term pathology. HSE can trigger autoimmune encephalitis, particularly anti-NMDAR encephalitis, which manifests with neuropsychiatric and cognitive symptoms resembling AD and amnestic mild cognitive impairment (aMCI) (55). Thus, recurrent viral activity and post-infectious autoimmunity may act in concert to drive progressive neurological dysfunction.

The immunosuppressed cohort provided further insights. Cyclophosphamide/dexamethasone-induced moderate clinical signs emerged within days, but by 20 dpi only low-level UL19 signals were detectable, while caspase-3 activity was elevated in both infected and control animals. This suggests that reactivation likely occurred shortly after treatment but was missed due to the timing of euthanasia. Glucocorticoid-induced depletion of T cells (56, 22) may have further obscured detection. These limitations highlight the need for shorter treatment-to-sampling intervals in future studies to better capture reactivation events.

In summary, our long-term study demonstrates that alphaherpesvirus latency is not confined to the TG but can be established in multiple CNS regions, with potential for sporadic reactivation and chronic low-grade inflammation. By integrating molecular, histopathological, and clinical readouts, we show how lytic activity and immune surveillance may contribute to long-term neuropathology. This model provides a platform for addressing unresolved questions, including whether episodic reactivation or chronic infection is the primary driver of CNS disease progression.

## Material and methods

### Virus

The attenuated PrV mutant PrV-ΔUL21/US3Δkin, derived from the PrV wildtype strain Kaplan (57) was previously described (29, 58). Virus stocks were propagated in rabbit kidney (RK13) cells maintained at 37°C in minimum essential medium (MEM) supplemented with 10% fetal calf serum (FCS) (Invitrogen).

### Animal experiments

Female CD1 mice (6-8 weeks old, Charles River Laboratories) were used as the standard infection model (29). Animals were housed in groups up to five in conventional cages (type II L) under Biosafety Level 2 (BSL 2) conditions at the experimental animal facility of the Friedrich-Loeffler-Institut, Greifswald-Insel Riems. Housing conditions included a 12h light/dark cycle (light intensity 60%), a temperature of 20-24°C, and ad libitum access to a standardized diet (ssniff Ratte/Maus-Haltung) and fresh drinking water. Bedding (ssniff Spezialdiäten Abedd Espen CLASSIC), nesting material (PLEXX sizzle nest), and environmental enrichment (PLEXX Aspen Bricks medium, mouse smart home, mouse tunnel) were provided.

After a 7-day acclimatization period, mice were deeply anesthetized by intraperitoneal injection of 200µl ketamine/xylazine (ketamine: 60 mg/kg; xylazine: 3 mg/kg, diluted in 0.9% NaCl). A total of 5 µl of virus suspension containing 1 × 10^4^ plaque forming units (PFU)/ml of PrV-ΔUL21/US3Δkin were administered intranasally per nostril. Mock-infected mice received cell culture supernatant from uninfected RK13 cells.

Study 1: Twelve groups (n = 7 per group) were inoculated either with PrV-ΔUL21/US3Δkin or mock solution. Animals were euthanized at 21, 42, and 105 dpi. Groups 1–6 were analyzed by RNAscope™ and histopathology; groups 7–12 were analyzed by RT-qPCR.

Study 2: As described previously (30), six groups (n=6 per group) infected with PrV-ΔUL21/US3Δkin and one group mock treated, were euthanized at 28, 35, 42, 49, 84 and 168 dpi. An additional group (n= 10), 5 animals infected with PrV-ΔUL21/US3Δkin and 5 mock treated, received an immunosuppression at 170 dpi via intravenous injection of 5 mg cyclophosphamide and 0.2 mg dexamethasone in 250 µL phosphate-buffered saline (PBS). These mice were sacrificed 20 days later for histopathological and RNAScope™ analyses.

Randomization was performed prior to the experiment to assign animals to treatment groups and time points. Clinical evaluation was continuously monitored (24/7) using a standardized scoring system (29) encompassing three categories: (I) external appearance, (II) behavior and activity, and (III) body weight.

Scores ranged from 0 to 3; animals reaching a score of 3 in any category or 2 across all three were euthanized to meet humane endpoint criteria. For euthanasia, animals were pre-treated with Carprofen (Rimadyl, Pfizer, 10 mg/kg, subcutaneous) analgesia, followed by deep isoflurane anesthesia. Transcardial perfusion was performed via the left ventricle with PBS and subsequently 4% paraformaldehyde (PFA) as described by (59). Mice were euthanized by decapitation at the level of the first cervical vertebra.

In study 1, brains from groups 1-6 were post-fixed in 4% PFA for at least one week, brains from groups 7-12 were sectioned into six regions (see section RT-qPCR) and stored in PBS for molecular analysis. In study 2, entire heads were fixed in 4% neutral-buffered formalin and decalcified in Formical 2000 (Decal, Tallman, N.Y.) for at least three days.

### Histopathological analysis

Brains from three PrV-ΔUL21/US3Δkin inoculated mice euthanized at 11-14 dpi (humane endpoint), 42 dpi, and 105 dpi (study 1), as well as heads from three infected mice euthanized at 28 dpi and from the cyclophosphamide/dexamethasone group (study 2) were processed for histopathology. Additionally, brains/heads from one mock-inoculated animal (study 1) and one mock-inoculated, immunosuppressed animal (study 2) served as controls.

Each brain/head was sectioned into six coronal levels from rostral to caudal, embedded in paraffin wax, and cut at 3 µm thick sections using a rotating microtome (Hyrax M55, Zeiss). Anatomical landmarks were defined according to Rao (60), resulting in six standardized levels: olfactory bulb (L1), prefrontal cortex (L2), frontoparietal cortex and basal ganglia (L3), parietal cortex, thalamus, hypothalamus, and hippocampus (L4), midbrain (L5), and cerebellum/pons (L6) (Fig. 3A, L1–L6). For light microscopy, sections were mounted on Super-Frost-Plus-Slides (Carl Roth GmbH, Germany) and stained with hematoxylin and eosin (H&E). Neuropathological analysis focused on CNS inflammation, neuronal necrosis and reactive gliosis.

Slides were examined using a Zeiss Axio Scope.A1 microscope equipped with 5x, 10x, 20x, and 40x N-ACHROPLAN objectives (Carl Zeiss Microscopy GmbH, Jena, Germany). Whole slide-scans were obtained with a NanoZoomer digital slide scanner (Hamamatsu, S60).

#### Immunohistochemistry

Immunohistochemistry was performed to identify infiltrating immune cells and apoptotic processes in relation to histopathological changes. Primary antibodies used are listed in Table 1.

**Table 1:** Primary antibodies used for immunohistochemistry.

| antigen | target | manufacturer | clonality/host species | working dilution |
| --- | --- | --- | --- | --- |
| Iba 1 | microglia/<br>macrophages | FUJIFILM Wako,<br>01-19741 | polyclonal rabbit | 1:500 |
| GFAP | astrocytes | Abcam, ab16997 | polyclonal rabbit | 1:400 |
| CD3 | t-cells | DAKO, A0452 | polyclonal rabbit | 1:100 |
| cleaved<br>caspase-3 | apoptosis | Cell Signaling<br>Technology,<br>9661 | polyclonal rabbit | 1:800 |

Paraffin-embedded tissue sections were dewaxed and rehydrated. Endogenous peroxidase activity was blocked with 3% hydrogen peroxide (Merck, Germany) for 10 min. Antigen retrieval was performed in 10mM citrate buffer (pH 6.0, without detergent) for Iba1, GFAP and Cas-3, or in 10mM Tris-EDTA buffer (10mM Tris base, 1mM EDTA solution, pH 9.0) for CD3. Sections were heated for 20 min in a pressure cooker (Sichler, Germany, NX-3213-675) for epitope demasking. After rinsing in Tris-buffered saline (TBS), nonspecific binding was blocked using normal goat serum (diluted 1:2 in TBS, 30min). Sections were incubated overnight at 4°C with primary antibodies diluted in TBS. The following day, slides were washed with TBS and incubated with biotinylated goat anti-rabbit IgG (Vector Laboratories, BA 1000, 1:200) for 30min at room temperature. For Iba1 and Cas-3 staining, detection used avidin-biotin-peroxidase (ABC) complex (Vectastain Elite, PK 6100), for 30 min at RT. For CD3 and GFAP, Polymere ImmPress®+ System (DAKO, MP-7451) was applied. Antigen-antibody complexes were visualized using AEC-substrate (abcam, ab64252), yielding red signal deposition. Slides were rinsed with deionized water, counterstained with Mayer’s Hematoxylin for 10 min, and coverslipped using Aquatex (Merck).

#### Scoring of inflammatory cells

Inflammatory responses were assessed at 11–14, 28, 42, 105, and 190 dpi in eight anatomically defined brain regions: OB, AI, Pir, LEnt, S1, Sp5, TG and Cb. Regions of interest (ROIs) were defined according to *The Mouse Brain in Stereotaxic Coordinates* (61). Each region was systematically evaluated in standardized coronal brain sections : OB (L1), AI (L2/3), Pir (L2/3/4), LEnt (L5), S1 (L2/3/4), Sp5 (L6), Cb (L6), TG (L3/4) (S2 Fig). For regions analyzed at multiple levels, the highest score per region was recorded.

CD3^+^ T-cell infiltration was semiquantitatively scored according to a published system (Table 2 (30)). Cas-3 was scored using the same criteria. Iba1⁺ microglia/macrophages and GFAP⁺ astrocytes were qualitatively assessed as present or absent (Table 3), based on glial activation regions with CD3⁺ infiltration. All evaluations were perfomed at high-power (20x or 40x).

**Table 2:** Semiquantitative scoring of CD3 and cleaved caspase-3 immunoreactivity in murine brain sections.

| score | CD3/ cleaved caspase-3 |
| --- | --- |
| 0 | absent |
| 1 | <10 cells |
| 2 | 11-20 cells |
| 3 | > 20 cells |

**Table 3:** Qualitative assessment of Iba1 and GFAP immunoreactivity in murine brain sections.

| score | Iba- 1/ GFAP |
| --- | --- |
| - | absence of glial activation |
| + | glial activation/ cluster |

The scoring system was applied to assess CD3⁺ T cell infiltration and cleaved caspase-3⁺ apoptotic cells across brain regions. Immunostaining for CD3 and cleaved caspase-3 was performed on coronal brain sections representing six anatomical levels. Selected regions of interest (ROIs) were evaluated; if multiple levels were available, the highest score per region was used. Scores ranged from 0 to 3 and were assigned according to the number of positive cells detected by light microscopy.

Immunostaining for Iba1 and GFAP was performed on coronal brain sections representing six anatomical levels. Regions of interest (ROIs) were analyzed at these levels; if multiple levels were available, the highest score per region was recorded. Glial activation was assessed qualitatively and categorized as present (+) or absent (−), based on the presence of immunoreactive cells with reactive morphology.

#### In situ Hybridization (RNAscope™)

Detection of PrV mRNA transcripts was performed on FFPE sections (Fig 3A) using the RNAscope™ 2.5 HD Duplex Reagent Kit (ACD. Inc, Cat. No. 322430) (35).

A C1 probe targeting the lytic viral transcript UL19 (V-SHSV-UL19, 66973–68498 base pairs; ACD Inc., Cat. No. 548251) was detected as green punctate signals via horseradish peroxidase (HRP)-mediated chromogenic reaction.

A C2 probe targeting the large-latency transcript (LLT) region of the latency-associated transcript (LAT) (V-SuHV1-LLT-O2-C2, 94664-95952 base pairs, ACD. Inc, Cat. No. 1808611) was detected as red punctate signals by alkaline phosphatase (AP)-mediated chromogenic reaction (Fig 3B). Probes were mixed at a dilution of 1:50 (C2 in C1).

As technical control, the murine housekeeping gene - Ppib (Mm-Ppib, ACD Inc., Cat. No. 313911, C1), encoding the peptidyl-prolyl cis-trans isomerase B, was combined with ubiquitin C (Mm-Ubc, ACD. Inc, Cat. No. 310779, C2) at a dilution of 1:50 (C2 in C1). An Escherichia coli DapB probe (Duplex Negative Control Probe, ACD. Inc, Cat. No. 320759) served as negative control (S3 Fig).

##### Pre-treatment and hybridization

Slides were baked at 60°C for 1 hour, deparaffinized in xylene (2×5 min), and rehydrated in 100% ethanol (2×1 min) at RT. After air-drying, endogenous peroxidase activity was quenched using hydrogen peroxide (H₂O₂; ACD Inc., Cat. No. 322330) for 10 min at RT. Target retrieval was performed by boiling slides in antigen retrieval buffer (ACD Inc., Cat. No. 322000) using a pressure cooker (Sichler, Germany, NX-3213-675) for 15 minutes. Slides were washed in distilled water, and dehydrated in 100% ethanol. After air-drying, sections were circumscribed with a hydrophobic barrier pen (Vector Laboratories, H-4000) to confine reagent application to the tissue area. Slides were then treated with Protease Plus (ACD Inc., Cat. No.322330) for 15 minutes at 40°C in the ACD HybEZ™ II Hybridization System (ACD Inc., Cat. No. 321711) (HybEZ oven). Slides were rinsed in distilled water and incubated with the specific probes and controls for 2 hours at 40 °C in the HybEZ oven, followed by two washes (2 minutes each) in the wash buffer (ACD Inc., Cat. No. 310091). Afterwards, the slides were stored overnight at RT in a 5X saline-sodium citrate buffer (SSC) (20X SSC, 1:4, diluted in distilled water; 20x SSC: 175 g of NaCl and 88 g of sodium citrate in 1 L distilled water, pH= 7.0).

##### Signal amplification and detection

On the following day, slides were washed twice in wash buffer and preamplification was performed by applying amplicon (Amp) 1 for 30 minutes at 40 °C. Further amplification steps were carried out sequentially with Amp 2 (15 min) and Amp 3 (30 min), both at 40 °C. Detection of C2 (LAT) was achieved with Amp 4 (15 min at 40°C) in the HybEZ oven, followed by Amp 5 (30 min at RT), and the AP blocker Amp 6 (15 min at RT). For visualization of the red signal, slides were treated with a 1:60 mix of Red-B and Red-A (10min at RT). Detection of C1 (UL19) signal was performed using Amp 7-10 (Amp 7: 15mins, 40°C; Amp 8: 30mins, 40°C: HRP enzyme-solution; Amp 9: 30mins, RT; Amp 10: 15mins, RT), followed by green chromogenic development using a 1:50 mix of Green-B to Green-A (1:50) (10mins at RT). Between each step (Amp: 1-10 and chromogen development) slides were washed twice in wash buffer for 2 mins. Slides were counterstained with Mayer’s Hematoxylin (30 s at RT), blued in tap water (5min), dried at 60°C (15-30min) on a heat plate (MEDITE, BD00575) and cooled for 5 min. Finally, slides were immersed in fresh xylene (5 min) and coverslipped using EcoMount (Biocare Medical, BRR897L).

#### Semi quantitative analysis of *In-Situ* Hybridization

For the semi-quantitative analysis of PrV mRNA expression, eight anatomically defined brain regions were analyzed as described in the section scoring of inflammatory cells. Each RNA transcript appeared as a discrete chromogenic dot within the tissue section (LAT: red; Ul19: green). Signals were assessed separately for UL19 and LAT in each predefined brain region. Scoring of positive cells was carried out as illustrated in Fig 5, based on the number and distribution of RNA-positive signals within the ROI. Briefly, punctate signals were scored on a scale from 0 to 3: score 0 represented no detectable signal, score 1 corresponded to 1–20 signals, score 2 to 20–100 signals, and score 3 indicated more than 100 signals. Clustered signals were evaluated separately using scores 4 and 5, with score 4 assigned to 1– 20 clusters and score 5 to more than 20 clusters.

### Real-time quantitative PCR (RT-qPCR)

Brains from six PrV-ΔUL21/US3Δkin- or mock-inoculated mice per time point (9-10, 21, 42, and 105 dpi, study 1) were dissected into six regions: OB, Pir, TL, Cb, BS and TG. Tissues were homogenized in PBS with 5mm steel-beads (Fabrikat Martin) using a bead beater (Retsch, MM200). Total RNA was extracted using the QIAamp Viral RNA Mini Kit (Quiagen, Cat. No. 52904), and RT-qPCR was performed to detect transcripts of UL19 (lytic gene) and the latency-associated transcript (LAT). Custom-designed primers (Eurofins) were used for LAT, and validated primer-probe sets for UL19 were kindly provided by Dr. Conrad Freuling, Friedrich-Loeffler-Institut, Greifswald-Insel Riems. Primer sequences are listed in Table 4. β-actin served as the internal housekeeping gene. A long LAT oligomer was used both as a standard for generating the calibration curve and as a positive control. The threshold cycle (Ct) cutoff was set at 40, based on dilution series of the LAT standard. Reactions above this threshold were considered negative.

**Table 4.**
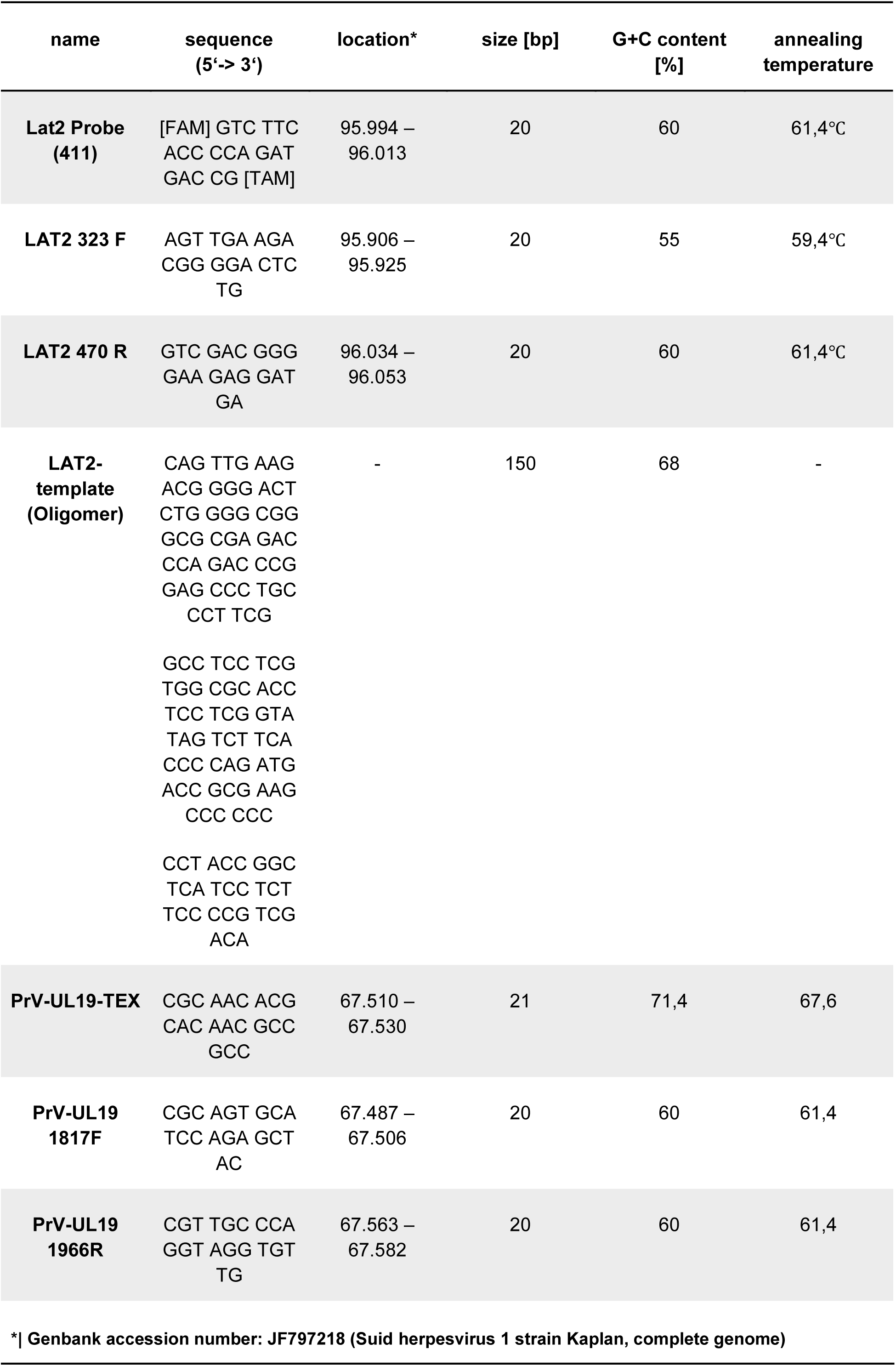
Primer sequences and characteristics for RT-qPCR detection of LAT and UL19 transcripts.

A 200µl primer-mix was prepared, containing 100pmol each forward and reverse primer, 2,5µl of the fluorophore-labelled probe, and 157,5µl of 0,1X TE buffer (pH 8,0). Each 20 μl PCR reaction consisted of:10 μl ready-to-use mix (2x SensiFast Probe No-ROX One-Step mix), 0,2 reverse transcriptase, 0,4 RiboSafe RNase inhibitor, 0,6 µl RNase-free water, 1,6µl of each primer-mix and 4µl RNA template. Amplification was performed using a Bio-Rad CFX96 thermal cycler under the following conditions: reverse transcriptase treatment at 45 °C for 15 min, inactivation of the transcriptase at 95 °C for 2 min, template denaturation at 95 °C for 15 sec, and annealing/amplification of the target cDNA at 60 °C for 40 sec (40 cycles). Fluorescence was measured during the elongation phase.

Primer pairs were designed for quantitative PCR analysis targeting latency-associated transcript (LAT) and the lytic viral gene UL19 of PrV-ΔUL21/US3Δkin. The table lists primer sequences, expected amplicon sizes, GC content and melting temperatures (Tm) used for specific amplification.

### Statistical analysis

Statistical analyses and graphical presentation were performed in GraphPad Prism 10.2.1 (GraphPad Software, Boston, USA).

Ct values obtained from RT-qPCR were analyzed by two-way analysis of variance (ANOVA) with Sidak’s multiple comparison test. Geometric mean values of UL19 and LAT transcripts were compared between brain regions. Correlations between CD3⁺ T-cell scores and UL19 RNA expression in corresponding brain regions were assessed using Spearman’s rank correlation coefficient.

p ≤ 0.05 was considered statistically significant and indicated by an asterisk in the figures.

## Acknowledgements

The authors would like to thank Silvia Schuparis and Robin Brandt for their excellent technical assistance. We are particularly grateful to Conrad Freuling and Thomas Müller for their valuable advice on RT-qPCR methodology. We further acknowledge Angele Breithaupt, Tobias Britzke and Lukas M. Michaely for scientific discussion.

## Ethics statement

Animal experiments were approved by the State Office for Agriculture, Food Safety and Fishery in Mecklenburg-Western Pomerania (LALFF M-V) with reference number 7221.3-1-034/ 22.

## Data availability statement

All data supporting the findings of this study are included in the Supporting Information files and are available from the corresponding author upon request, in compliance with PLOS’ data policy.

## Supporting information

**S1 Fig. Clinical condition of PrV-infected mice at euthanasia and corresponding heatmaps of UL19 and LAT gene expression across defined brain regions.** (A) The clinical condition of each mouse at the time of euthanasia is shown in relation to corresponding gene expression. Mice marked with an asterisk (*) exhibited severe clinical signs and reached predefined humane endpoint criteria, necessitating euthanasia. (B, C) RT-qPCR was performed to quantify RNA transcripts of UL19 (B) and LAT (C) of PrV in selected brain regions (OB = olfactory bulb, Pir = piriform cortex, TL = temporal lobe, TG = trigeminal ganglion, Cb = cerebellum, BS = brainstem) at defined time points. Heatmaps were generated from Ct values to illustrate spatial and temporal patterns of viral gene expression. Lower Ct values (darker color intensity) indicate higher transcript abundance.

**S2 Fig. Delineation of brain regions for semiquantitative analysis of immunohistochemistry and in situ hybridization.** Anatomical boundaries of each brain region were defined according to The Mouse Brain in Stereotaxic Coordinates (Paxinos & Franklin, 2001). Colored overlays indicate the specific areas analyzed within coronal sections at levels 1-6. Yellow: olfactory bulb (OB); light green: primary somatosensory cortex (S1); orange: agranular insular cortex (AI); blue: piriform cortex (Pir); grey: trigeminal ganglion (TG); pink: hippocampus (Hpc); purple: lateral entorhinal cortex (LEnt); brown: cerebellum (Cb); dark green: spinal trigeminal nucleus (Sp5). Brain atlas images adapted from The Mouse Brain in Stereotaxic Coordinates (Paxinos & Franklin, 2001).

**S3 Fig. Representative images of RNAscope™ detection in the murine brain**. (A) Negative control probe targeting Escherichia coli DapB shows absence of specific signals, confirming assay specificity. (B) Positive control probes for murine Ppib (C1, green) and Ubc (C2, red) demonstrate robust and widespread expression in neuronal tissue, validating RNA integrity and hybridization efficiency. Scale bar = 250 µm.

**S1 Table. Summary of clinical findings, histopathological diagnoses, and molecular analyses**. Each mouse is listed with clinical signs, HE-based pathological evaluation, and corresponding scores from IHC and RNAscope™ (ISH). Animals that reached the humane endpoint are indicated with an asterisk (*). TG = trigeminal ganglion, Cb = cerebellum, Sp5 = spinal trigeminal nucleus, S1 = primary somatosensory cortex, Hpc = hippocampus, LEnt = lateral entorhinal cortex, Pir = piriform cortex, AI = agranular insular cortex, OB = olfactory bulb.

